# Molecular dynamics study of phospholipid membrane electroporation induced by bipolar pulses with different intervals

**DOI:** 10.1101/2022.07.21.501059

**Authors:** Fei Guo, Jiong Zhou, Ji Wang, Kun Qian, Hongchun Qu

## Abstract

In this study, PM-EP induced by bipolar pulses with different intervals was investigated by all-atom molecular dynamics simulation. Firstly, PM-EP was formed during the positive pulse of 2ns and 0.5V/nm, then the effects of various intervals of 0, 1, 5, 10ns on PM-EP evolution were investigated, and the dynamic changes of different degrees of PM-EP induced by following negative pulses of 2ns and 0.5V/nm were analyzed. The elimination of the PM-EP during the interval of bipolar pulses were determined and it was related to the degrees of PM-EP and the time of intervals, then the degrees of PM-EP at the end of the intervals were classified and quantitatively defined, namely, Resealing, Destabilizing and Retaining state. These three states appeared due to the combined effect of both the preceding positive pulse and the interval. Furthermore, the evolution of PM-EP in resealing state under negative pulses was similar to that of positive pulses as evidenced by EP formation time and degree of PM-EP, the destabilizing state had the same trends as the resealing state except that the re-electroporation of phospholipid membrane appeared faster and larger degree of EP obtained with the same pulse exposure time. Regarding the retaining state, the negative pulses enhanced PM-EP with more profound water bridges, which can be considered as the effect of electric field superposition. These results can improve our understanding of the fundamental mechanism of bipolar pulse-induced PM-EP.

**Graphical Abstract:** 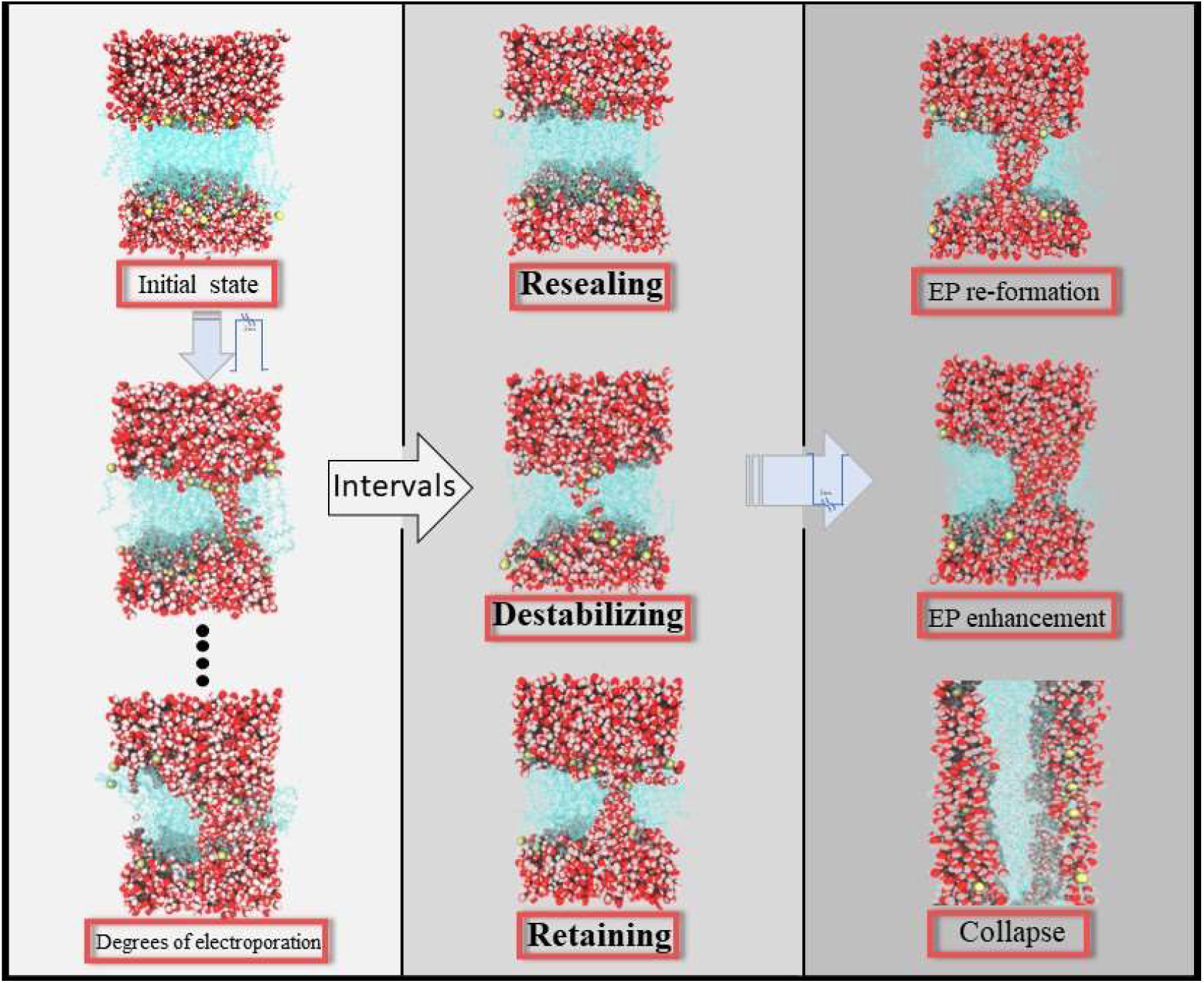

**Highlights:**

- Quantitative and qualitative definition of the three states of the phospholipid membrane electroporation at the end of the intervals.
- Clarification that the states of phospholipid membrane electroporation were generated due to the combined effect of positive pulses and intervals.
- Quantitative and qualitative comparison the evolution of the states of phospholipid membrane electroporation during the negative pulses.

## 1. Introduction

The permeability of cell membranes increases when exposed to a continuous external electric field, allowing extrinsic substances such as anticancer drug that were normally blocked outside the membrane to enter, and the process is known as EP [1], and has been widely used in medical and biological fields [2].

Multiple studies have demonstrated that the bipolar pulses (BP) have important roles in EP areas such as changing the permeabilization of cells [3], cellular mortality [4], and reducing muscle contraction [5]. The application of BP produced a physiological phenomenon called bipolar pulse cancellation (BPC) [6], which refers to the reduction or elimination of the permeability of cell membranes when the system was subjected to BP compare with the corresponding unipolar pulses. Another phenomenon that is opposite of BPC and called bipolar pulse enhancement (BPE), which refers to the increased or enhanced of the permeability of cell membranes affected by BP. Researchers have found that the intervals can influence the result of BP [7]. For example, Polaj*ž*er et al. implemented eight bursts of 1–10 ms pulses with interval delays of 0.5 ms–10 ms and estimated the permeability of phospholipid membrane and cellular mortality, they revealed that interval of pulses can eliminate BPC [8]. Valdez et al. surveyed different intervals between symmetrical and asymmetrical nanosecond electric pulses to detect their influence and discovered that the BPC could be eliminated by changing the interval of the pulses, even achieving the same effect as the unipolar pulses [9], which produces similar EP and membrane damage as unipolar pulses [10], that the duration and interval of the pulses together determine whether the BPC or BPE exists. So far, the exact mechanism of interval of BP on cell EP is still unclear, and further theoretical and/or in silicon studies are required to address this issue at the atomic level.

Molecular dynamics (MD) is a thermodynamic calculation method based on Newtonian deterministic, which can provide microstructure information of higher spatial resolution and structural dynamics information of higher temporal resolution simultaneously, which was an effective tool to investigate the molecular mechanism of PM-EP [11][12]. Vernier et al. tracked with FM1-43 fluorescence and showed that in contrast with unipolar pulses, BP redistribute phospholipids at both the anode and cathode side of the cell [13]. Tang et al. compared EP formation time by bipolar and unipolar pulses using MD simulation and found that the appearance of BPC or BPE depends on pulse width and intensity [14]. S*ö*zer et al. used systematic variations in initial MD simulation conditions to measure the lifetime of lipid bilayer EP and found that the kinetics of ultrashort electric field-induced giant unilamellar vesicles (GUV) permeabilization differed from the disclosed results in cells exposed to ultrashort electric fields [15]. Brand et al. obtained the RMS fluctuations of the amide proton using MD simulation and provided bilinear model along with the NH coupling constant [16]. Although MD simulation has been applied to many studies, very few studies have systematically investigated the effect of BP interval on EP process at the molecular level.

In this study, PM-EP induced by BP with various intervals was investigated by all-atom molecular dynamics simulation. Firstly, positive pulses of 2ns and 0.5V/nm were applied to the phospholipid membrane to produce EP, then the effects of 0, 1, 5, 10ns intervals on evolution of PM-EP were analyzed, and the dynamic changes of various degrees of PM-EP induced by negative pulses were also analyzed. The PM-EP were analyzed using parameters such as interfacial water average dipole moment (IDM), time of EP formation under the positive pulses (*t_ep_*), the number of phospholipid heads between the phosphorus layers (*P_N_*),the number of water molecules between the phosphorus layers (*W_N_*) and the time of EP regeneration under negative pulses (*t_rep_*).

## 2. Materials and methods

### 2.1. Model system

The simulations were performed using a phospholipid membrane model of 6.0 nm × 6.0 nm × 8.2 nm [17]. The phospholipid membrane was wrapped in a 0.4 mol/L KCl solution and uniformly consisted of 108 1-palmitoyl-2-oleoylsn-glycero-3-phosphocholine (POPC) lipid molecules which are important components of organelle membranes and cell membranes [18], there are 4912 water molecules in the model, as shown in figure supplement S1.

### 2.2. Simulation parameters

All-atom MD simulations were carried out with the NAMD software package [19] and CHARMM27 force field [20] for simulation experiments and observed by VMD software [21]. Using the Langevin piston Nose-Hoover method, three-dimensional periodic boundary conditions were applied at constant temperature (310K) [22] and constant pressure (1atm) [23]. The velocity Verlet integration scheme was combined with the equation of motion [24], and a multi-time step algorithm with time step of 2fs was used [25]. Setting all bonds connected to the hydrogen atoms to be rigid [26]. Using the transition function to correct the introduction of van der Waals forces and short-range electrostatics, which starting at 1 nm and cutting off at 1.2 nm, using the PME algorithm to calculate the long-range electrostatic potential, setting the PME mesh size to 1 [27].

The simulation process was divided into two steps. The first is balanced, the model is generated in CHARMM-GUI and initial equilibrium is achieved by a stepwise release of bound phospholipids [28], then an all-atom MD simulation of 10ns was initiated to equilibrate the model system without an applied electric field, which includes multiple model energy minimization and energy balanced [29]. Next, the model is subjected to an external bipolar multi-interval pulse in the z-axis direction, as shown in Fig. 1. The pulses were divided into three parts: positive pulses of 2ns and 0.5V/cm, various intervals of 0, 1, 5,10 ns, followed by negative pulses of the same duration and intensity of the positive pulses.

**Figure 1:**
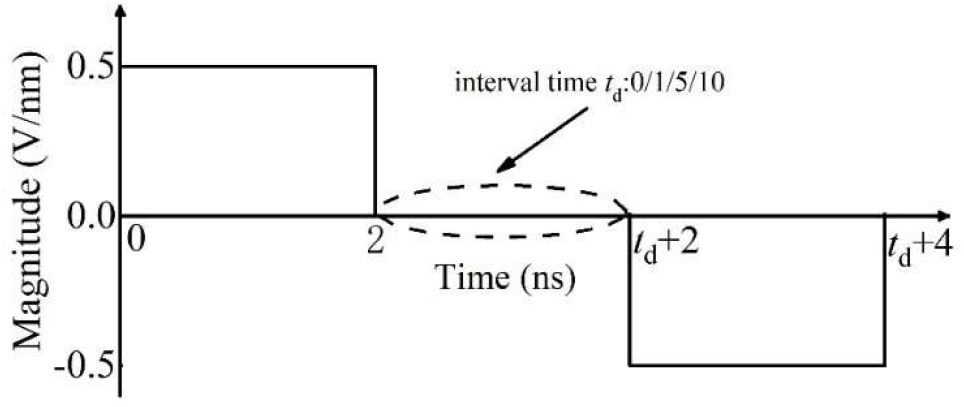
The positive pulses of 2ns and 0.5V/cm, various intervals of 0, 1, 5,10 ns, followed by negative pulses of the same duration and intensity of the positive pulses.

### 2.3. Basic Theory

The quantum mechanical equations were used in molecular dynamics for the evolution of the trajectories and states of motion of all atoms in the model. Therefore, the atomic motion of molecular dynamics was consistent with the fundamental laws of physics and can be represented by the Newtonian equations of motion as Eq. (1)

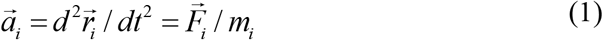

where 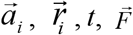 and *m_i_* represents acceleration, displacement, time, force, and mass of *i* atom.

The magnitude of 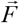 can be derived by finding the gradient of the potential function *U* as Eq. (2)

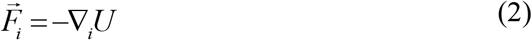

where 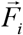, ∇_*i*_, *U* represents the magnitude, gradient operation, potential energy function. The potential energy function consists of interatomic bond energy, interatomic dihedral energy, atomic pinch energy, and Van der Waals interactions with electrostatic interactions.

## 3. Results

In this study, BP with various intervals were applied to the phospholipid membrane, each simulation had produced EP after applying the positive pulses and repeated 12 times. We observed the radius of the water bridges between phospholipid layers which could reflect the degrees of PM-EP at the end of the intervals in all simulations, and found that the PM-EP were qualitatively divided in three states:

Resealing: the water bridges completely disappeared at the end of the intervals as shown in Fig. 2(a).
Destabilizing: the water bridges changed to the water protrusions at the end of the intervals as shown in Fig. 2(a).
Retaining: the water bridges persisted at the end of the interval as shown in Fig. 2(a).

**Figure 2:**
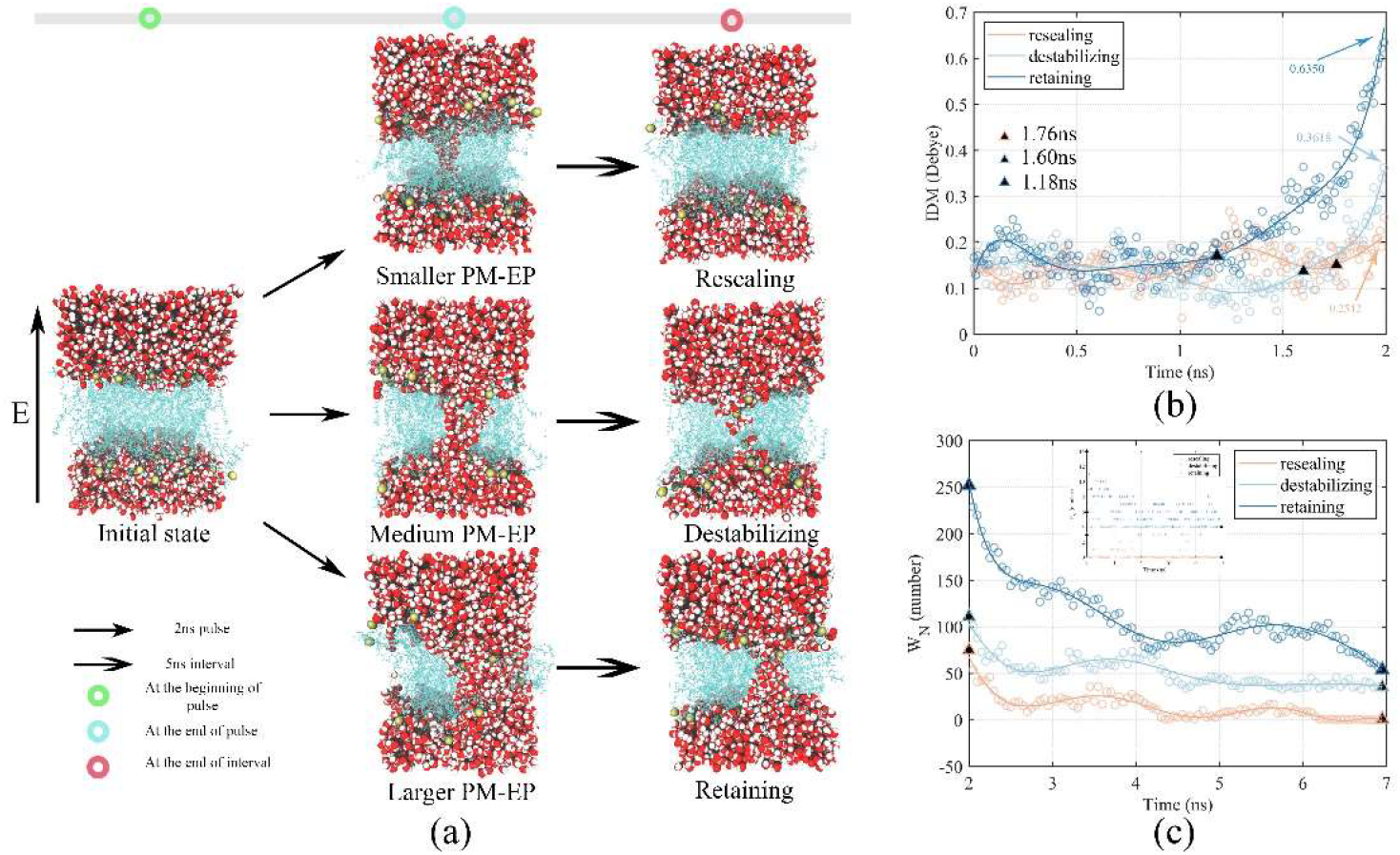
(a) phospholipid membranes changed into three states under the action of positive pulses of 2ns and 0.5V/nm and 5ns interval. White is oxygen atom, red is hydrogen atom, one oxygen atom and two hydrogen atoms form a water molecule, cyan indicates phospholipid molecule, brown indicates phospholipid head atom. (b) The time course of IDM during the positive pulses corresponding to the simulations in (a). The triangles on the curve indicates *t_ep_* in three states, and the value in the figure indicates the IDM at the end of the positive pulse. (c) *W_N_* and *P_N_* variation during the intervals corresponding to the simulations in (a). The triangles represent the beginning and end of the intervals, since the end values of *P_N_* in destabilizing state and retaining state are same represented by the same triangle.

The quantitative analysis of *P_N_*, *W_N_* and *t_rep_* in all simulations was in table supplement S1-S3. Accordingly, we found that the above three states conformed the rules as in Tab. 1.

**Table 1:** Parameters of states conformed the rules.

| State | $P_N$<br>(number) | $W_N$<br>(number) | $t_{rep}$<br>(ns) |
| --- | --- | --- | --- |
| Resealing | 0 | 0~16 | 0.50~2.00 |
| Destabilizing | 1~4 | 14~56 | 0.00~0.50 |
| Retaining | 4~16 | 52~178 | - |

Note: The water bridge of retaining state was always present and its $t_{rep}$ parameter was not counted.

### 3.1. Differential generation of PM-EP by positive pulses

Although the initial state of the phospholipid membrane was the same, the PM-EP did have differential states at the end of the intervals, as shown in Fig. 2(a). The same positive pulses were applied to the phospholipid membrane, there were considerable differences in PM-EP at the end of pulse which can be classified as smaller, medium, and larger as in Fig. 2(a). The smaller degree of PM-EP at the end of the positive pulses became resealing state after the 5ns interval. The medium degree of PM-EP in term of the radius of the water bridge, then changed into the destabilizing state. The larger degree of PM-EP changed into the retaining state.

We calculated the IDM of phospholipid membrane of the independent simulations corresponding to the three degrees of PM-EP in Fig. 2(a) when positive pulses were applied, as in Fig. 2(b). The *t_ep_* of resealing state was larger than others. Moreover, radiuses of the water bridges were positively correlation to the IDM at the end of the positive pulses. We then statistically investigated the relationship between indicators of PM-EP (IDM, *W_N_* and *P_N_*) at the end of the positive pulses and *t_ep_* with all simulations. The figure supplement S2 show a negatively correlation between indicators of PM-EP and *t_ep_*, which reflected the difference in *t_ep_* led to the various degrees of PM-EP at the end of the positive pulses. So, the different *t_ep_* could affect the states at the beginning of the intervals. *P_N_* was only used as an auxiliary reference quantity because the values were too small and more susceptible to random fluctuations.

Since the randomness of the EP phenomenon generated by the phospholipid membrane during the positive pulses, the degrees of PM-EP obtained at the end of the positive pulse were different. Smaller *t_ep_* indicated larger water bridges and IDM at the end of the positive pulses.

### 3.2. Elimination of PM-EP by intervals

In Fig.2(c), the maximum and minimum *W_N_* were obtained for the phospholipid membrane in the retaining and resealing state, respectively. In addition, biggest and smallest decreased in *W_N_* can be found in the retaining and resealing state during the 5ns interval. Therefore, the amount of *W_N_* and its decreased was positively correlation to the degree of PM-EP with the same interval.

The repetition number of the three states after applying four different intervals of the BPs was shown in Fig.3. The results of phospholipid membrane in 12 repeated simulations were all in retaining state with 0 ns interval. Phospholipid membrane was mainly categorized into the two states of destabilizing and retaining with 1ns interval. Three states of phospholipid membrane coexisted with 5ns and 10ns intervals, decreased in number of retaining state and resealing state increased can be found with the increasing interval, which meant that the elimination effect of intervals on the PM-EP was positively correlation to time. For example, the 1ns interval was too short to make the water bridges disappear completely, so the resealing state was not presented. When the interval was extended to 5 ns or 10 ns, the elimination effect of the interval on the PM-EP accumulated to the point where the water bridges completely disappeared and resealing state appeared.

**Figure 3:**
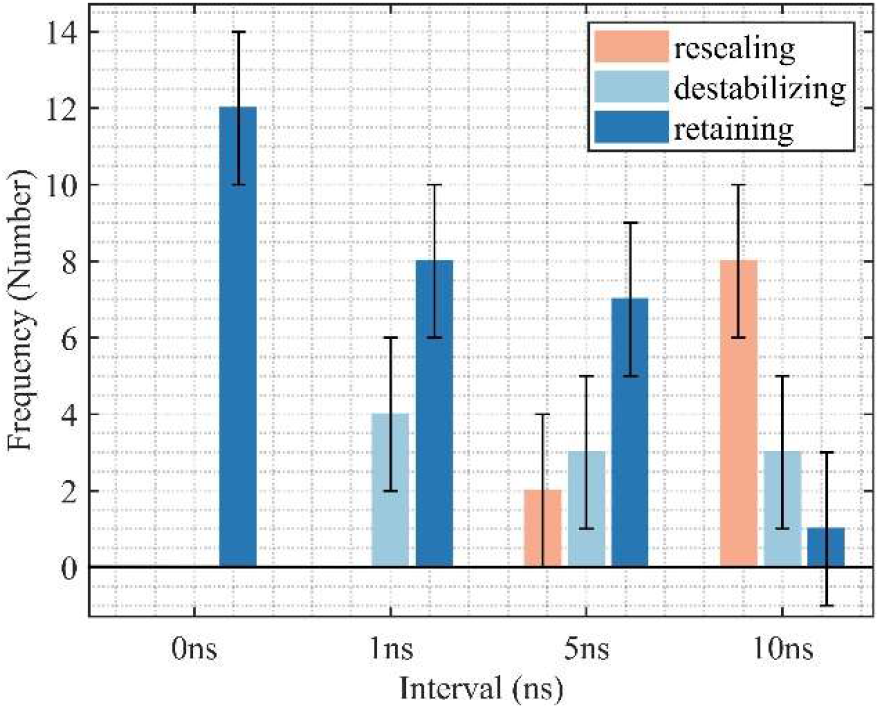
The repetition number of the three states at different intervals.

The progressively increasing of phospholipid membrane recovery in Fig. 4 (a) also demonstrated the cumulative elimination effect of intervals on PM-EP. The lipid tail was significantly bent when the phospholipid membrane was in the state that intervals lasting 0 ns. At 1 ns, the radius of the water bridges reduced and the deformed lipid tail gradually recovered. At 5 ns, the water bridges had broken into water protrusions and the lipid tail was further restored. At 10ns, water bridges had completely disappeared and the lipid tail leveled again. We analyzed the time course of *W_N_, P_N_* during the 10ns interval of Fig. 4(a), as shown in Fig. 4(b). *W_N_* and *P_N_* gradually decreased during the interval, which quantitatively demonstrated that the elimination of PM-EP was cumulative over time. Time dynamics of the average *W_N_* and *P_N_* with 10ns interval for each state was shown in Fig. 4(c). In retaining state, *W_N_* had a sharp decline at the beginning of the intervals of about 2ns, then linearly decrease for about 2ns, finally level off until the 10ns interval. Similar trends can be found in the destabilizing and resealing state but with smaller decreased slope at the beginning of the interval, then linearly decreased in *W_N_* until the end of the 10ns interval was observed in the resealing state. After fitting the data function of Fig. 4(c), the time courses of *W_N_* and *P_N_* in these three states during the 10 ns interval were consistent with the exponential change as in table supplement S4.

**Figure 4:**
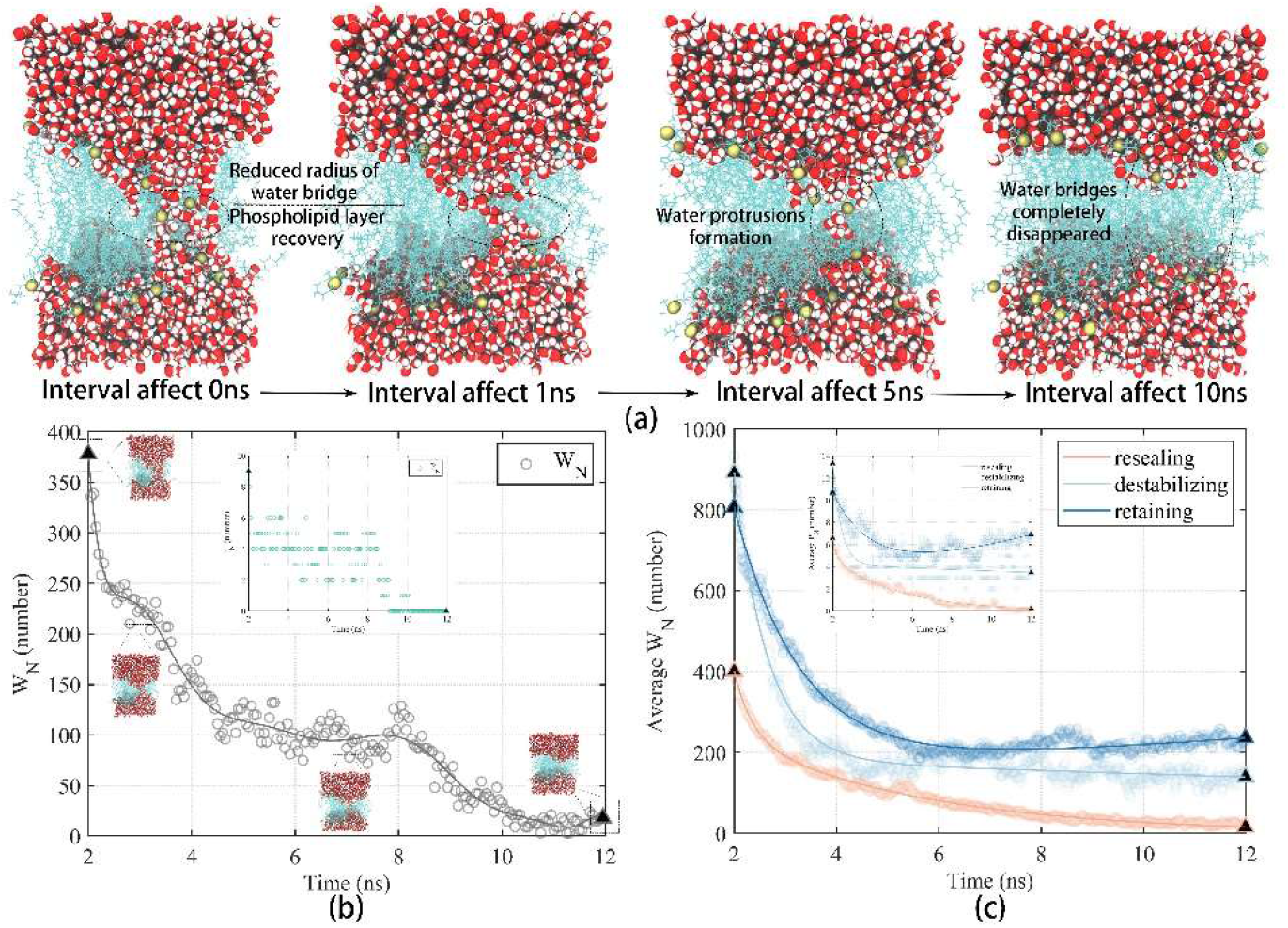
(a) The interval affects the PM-EP after the positive pulses. (b) *W_N_* and *P_N_* variation of (a) during the 10ns interval. (c) average *W_N_* and average *P_N_* variation for the three states with 10ns interval.

### 3.3. Evolution of the three states during the negative pulses

The states reflected the degrees of PM-EP at the end of the interval, and the trends in phospholipid membrane EP after applying negative pulses varied according to states. The resealing state with the negative pulses applied was shown in Fig. 5. The electric field was top-down generated inside the model, water molecules moved upward and away from the lower interface by the promotion of the electric field, gradually reaching in the water-lipid interface to form the water bridges [30][31]. There were no interfacial water molecules between the phospholipid layers as shown in Fig. 5(a) when the negative pulses were initially applied, so the IDM of the entire phospholipid layer was 0. At 0.7ns, water protrusions formed between the phospholipid layers as shown in Fig. 5(b,e), the portion of the phospholipid layers near the interface had non-zero IDM. At 1.5ns, the water bridge was stable and the IDM between the phospholipid layers became negative as shown in Fig. 5(c,f).

**Figure 5:**
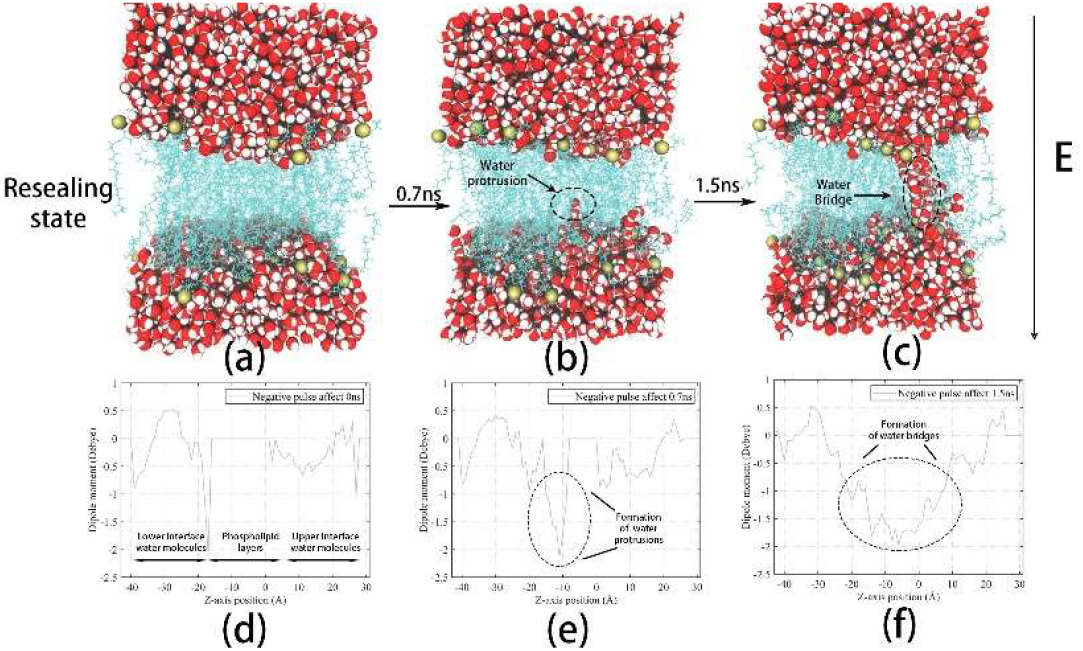
(a) 0ns, (b) 0.7ns, (c) 1.5ns after applying negative pulse to the phospholipid membrane in resealing state. (a)-(c) is the state at different time after applying negative pulses. (d)-(f) is the spatial location distribution of the phospholipid membrane along the Z-axis of the IDM.

Due to the upward movement of water molecules, the length of water protrusions at the upper interface (WPUI) decreased slowly, while the water protrusions at the lower interface (WPLI) increased continuously. Once the length of the WPUI was sufficient, the WPLI would quickly connect with the WPUI to form water bridges as shown in Fig. 6, if the length of the WPUI was not enough, the length would decrease continuously until it disappeared, and WPLI increased into the water bridges. The water bridges in the retaining state always existed when the negative pulse was applied, so the IDM between the phospholipid layers were always presented and increased with the continuous application of negative pulses as shown in Fig. 7. The evolution of PM-EP in retaining state with the negative pulses could be considered as the superposition of positive and negative pulses.

**Figure 6:**
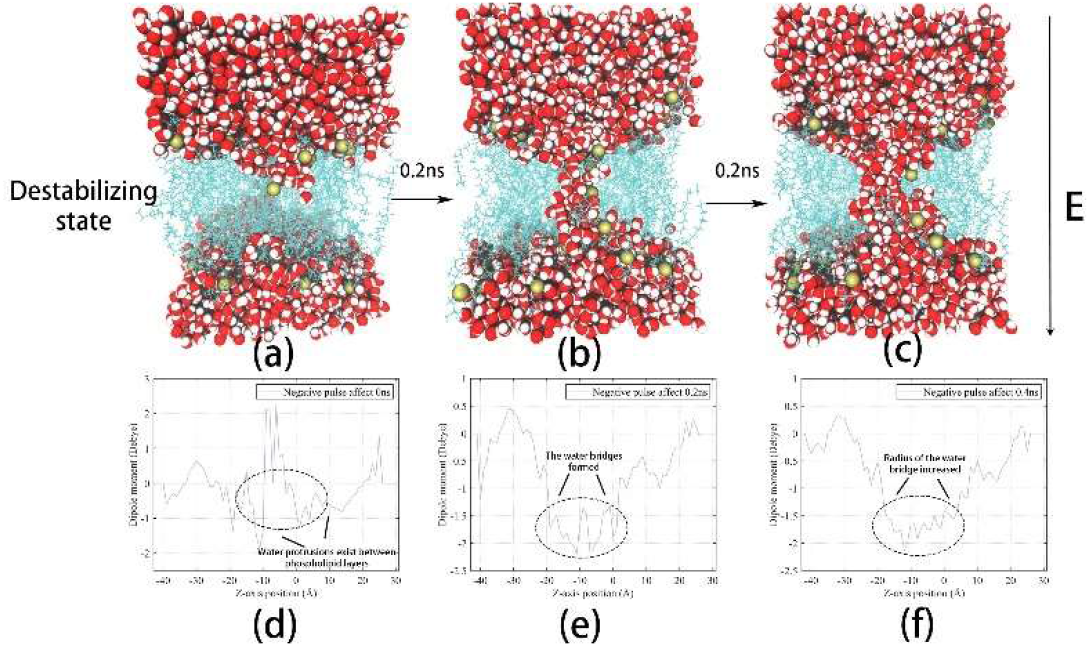
(a) 0ns, (b) 0.2ns, (c) 0.4ns after applying negative pulse to the phospholipid membrane in destabilizing state.

**Figure 7:**
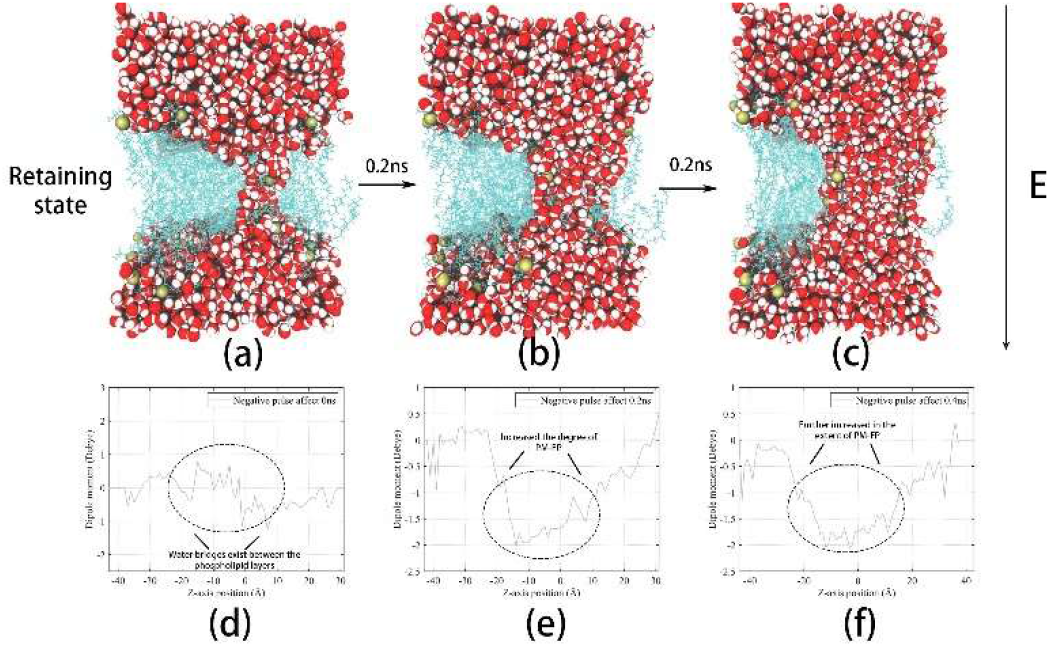
0ns, (b) 0.2ns, (c) 0.4ns after applying negative pulse to the phospholipid membrane in retaining state.

From Fig. 8 and Tab. 2, initial IDM and its rising rate in various states were different when the negative pulse was applied. Specifically, the degree of PM-EP and the rising rate of the IDM in the resealing state was the smallest. The IDM in destabilizing state raised faster than the resealing state due to the existence of the water protrusions between the phospholipid layers, next to the retaining state where water bridge always existed. We summarized 12 simulations at different intervals, the *t_ep_* and *t_rep_* in the destabilizing state was smaller than that in the resealing state as shown in table supplement S1-S3. Moreover, the *t_ep_* under the positive pulses of destabilizing state was greater than that of retaining state. So, we could think of destabilizing state as temporary state, which macroscopically manifested as water protrusions in the phospholipid layers and formed water bridges almost instantaneously after applying electric fields.

**Figure 8:**
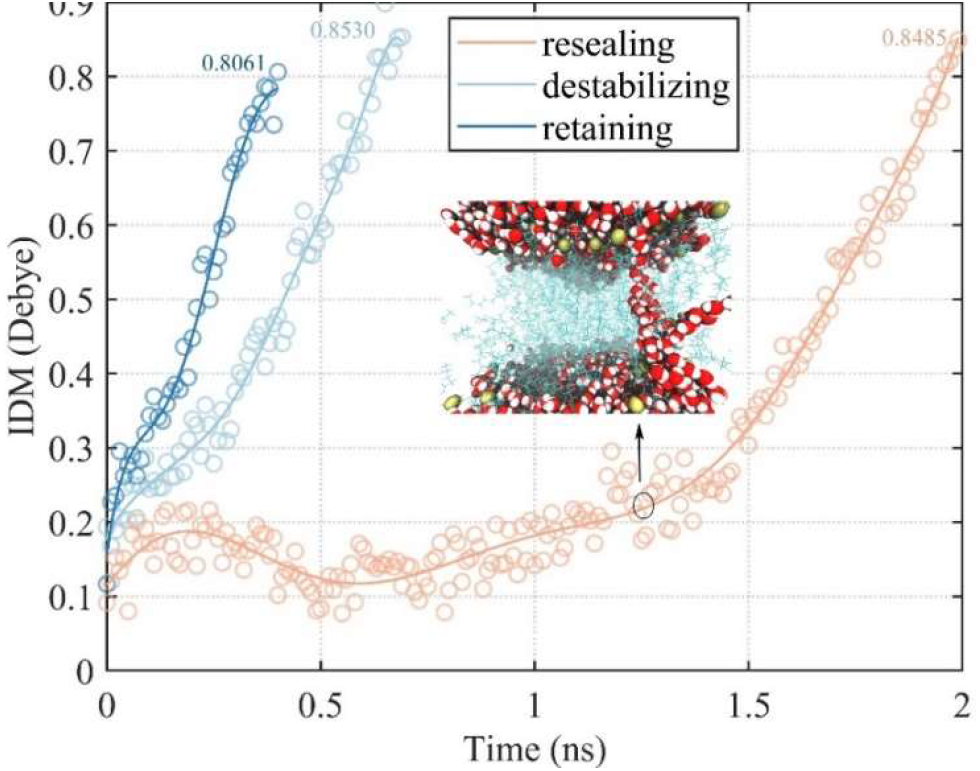
The absolute values of the IDM changes after applying negative pulse to the three states, the time is not the same because the model with too large degree of EP will collapse.

**Table 2:** Parameters of states after applying negative pulses

| State | Average IDM<br>(debye) | $t_{rep}$<br>(ns) | Repetition<br>frequency<br>(number) |
| --- | --- | --- | --- |
| Resealing | 0.1044 | 1.5990/6 | 10 |
| Destabilizing | 0.1354 | 0.2213/8 | 8 |
| Retaining | 0.1430 | - | 18 |

Note: Only 6 simulation data in resealing state are counted due to there is no EP formed in 4 simulations after applying negative pulses. The average IDM is the absolute value of the average IDM for states at the beginning of the negative pulses. The water bridge in retaining state has always existed and the results of phospholipid membrane in 12 repeated simulations were all in retaining state with 0 ns interval, so for not counting.

## 4. Discussion

In our study, the pulses are bipolar pulses with intervals of 0, 1, 5, and 10 ns and pulse width of 2 ns. The three states which represent the degrees of PM-EP at the end of the intervals were proposed quantitatively and qualitatively, and we found that the combined effect of the positive pulse and interval produced the three different states. The elimination effect of intervals on PM-EP was evaluated, and it was related to the degrees of PM-EP and the time of intervals. Finally, we compared the trends of PM-EP after applying negative pulse for the three states.

### 4.1. The exploration of interval

The interval of BP in changing the EP efficiency, regulating the BPC and BPE has been extensively studied [32][33], which also has a significant role in high-frequency biphasic pulses (HF-IRE) treatment outcomes and may improve the effectiveness of HF-IRE as clinical tissue ablation modality [34][35] [36][37] [38]. In this study, we varied the intervals within BP to observe the PM-EP dynamic at molecular level. The elimination effect of the interval on PM-EP was confirmed in all intervals [39][40], and cumulated over time [41]. This conclusion was in agreement with the results of Demiryure et al. who confirmed that the molecular transferred efficiency decreased monotonically with the delay time between pulses [42]. It is contrary to the concept of irreversible EP where the degree of EP was too large to recover that the elimination effect of interval was positively to the degree of PM-EP [43][44]. Saulis divided the process of pore resealing into three stages: pore reduction rapidly, pore reduction slowly, and pore closure completely, we then quantitatively described all states through *W_N_* and *P_N_* including the resealing state [45]. Inspired by Zachary et al. who used to divide the water bridge during the interval into four stages through *P_N_* [46], we divided the different degrees of PM-EP at the end of the interval into three states through *P_N_*, *W_N_*, and *t_rep_*.

### 4.2. Differentiation of states

The three states were defined in quantitative and qualitative perspectives. The resealing state is similar to the recovery of PM-EP [47][48]. If the interval lasts long enough, the retaining state is similar to irreversible EP [49]. The destabilizing state is the intermediate state between the resealing state and the retaining state. BPC were defined in previous study by tracking the amount of tracer entering the cell and whether the second part of the bipolar pulse reduced the influx of tracer [50]. There were many explanations for BPC, such as that BPC was balanced by local charging and discharging events across the membrane [9], where the effect of the first pulse was reduced by the second pulse of opposite polarity in the cancelled effect [34]. Since degree of EP reflected cell membrane permeability, the completely disappearance of the water bridge was consistent with the definition of BPC [51]. The resealing state of phospholipid membrane at the end of the intervals may correspond to the BPC, and the retaining state may correspond to BPE. The destabilizing state may be located in the intermediate state between BPC and BPE. As the destabilizing state changed in the negative pulse, BPE occurred if the degree of PM-EP was large enough when negative pulses was applied, and the opposite of generating BPC, this may provide some support for the mechanism of BPC and BPE generation.

### 4.3. Analysis of Simulation Limitation

The aim of this study was to investigate the effect of BP with intervals on the PM-EP, some conclusions were drawn from the results but there are still some limitations. First, our study investigated the effect of intervals on the PM-EP under BP, the number of repetitions for each interval is 12, so there were overall 48 simulations in this study. More simulations will greatly enhance the reliability of the simulation, but the current number of simulations is sufficient to reflect the relevant rules. In addition, the effect of parameters of the pulse such as whether the pulse is symmetric, the amplitude and the waveform of pulses on the frequency of the states occurring in the intervals will be studied subsequently. Second, several studies had measured the effect of various membrane components on PM-EP, such as the various types of phospholipids [52], the different levels of cholesterol [53], the ion channel proteins contained in the membrane [54]. It can be hypothesized that these components could affect the behavior of phospholipid membranes in the intervals. Specifically, whether *W_N_*, *P_N_* still exponentially changes during the intervals and the elimination effect of the interval on EP is enhanced or diminished, all need to be in-depth investigated in our future study. There are other parameters such as model dimensions, water-lipid ratios that affect the EP, which worth further exploration. Finally, most of the EP in the simulations were reversible, and irreversible EP had not been systemically studied. Despite the above limitations, this study analyzed the EP from the perspective of BP intervals, it paved the way for future research into the medical applications of BP as well as mechanistic studies still has a strong sense of guidance.

## 5. Conclusion

In this study, we applied BP with various intervals to the phospholipid membrane. The EP dynamics under the applied pulses were observed at the molecular level. We classified PM-EP at the end of the intervals into three states depending on the *P_N_*, *W_N_*, *t_rep_* and determined the action mechanism of positive pulses and intervals for the three states, as well as the tendency of the states changed during the intervals. We quantify the PM-EP evolution of the states during the following negative pulses. Our results confirmed the elimination effect of the interval on PM-EP and provided new guidance for the reperforation of PM-EP, and provide a new theoretical basis for further applications of EP technology in various fields.

## Acknowledgements

This work was supported in part by the Science and Technology Research Program of Chongqing Municipal Education Commission (Grant No. KJQN202100607), in part by the Natural Science Foundation of Chongqing, China (cstc2020jcyj-msxmX0393), and in part by the National Natural Science Foundation of China (No.51507024).

